# iSIM-sigma: efficient standard deviation calculation for molecular similarity

**DOI:** 10.1101/2024.11.24.625084

**Authors:** Kenneth Lopez Perez, Bill Zhao, Ramon Alain Miranda Quintana

## Abstract

The average and variance of the molecular similarities in a set is high-value and useful information for cheminformatics tasks like chemical space exploration and subset selection. However, the calculation of the variance of the complete similarity matrix has a quadratic complexity, *O*(*N*^2^). As the sizes of molecular libraries constantly increase, this pairwise approach is unfeasible. In this work, we present an alternative to obtaining the exact standard deviation of the molecular similarities in a set (with *N* molecules and *M* features) for the Russell-Rao (RR) and Sokal-Michener (SM) similarity indexes in *O*(*N M*^2^) complexity. Additionally, we present a highly accurate approximation with linear complexity, *O*(*N*), based on the sampling of representative molecules from the set. The proposed approximation can be extended to other similarity indexes, including the popular Jaccard-Tanimoto (JT). With only the sampling of 50 molecules, the proposed method can estimate the standard deviation of the similarities in a set with RMSE lower than 0.01 for sets of up to 50,000 molecules. In comparison, random sampling does not warrant a good approximation as shown in our results.

## Introduction

Binary fingerprints are one of the most common molecular representations in cheminformatics, they consist of a vector of ones and zeros.^1,2^ Each bit feature encodes structural information about the molecule, in the simplest case the presence or absence of a functional group/substructure.^2,3^ There are several other popularized fingerprint encoding methods; some are based in paths,^4^ topological,^5,6^ and radial information.^7,8^ One of the most popular applications of binary fingerprints is molecular similarity. ^1,2^ The importance of similarity lies in the property similarity principle, ”similar molecules have similar properties”,^9^ which is highly used in the rational drug design.^10,11^

There are several similarity indexes to quantify the similarity between molecules, for example, Russell-Rao (RR),^12^ Sokal-Michener (SM),^13^ Jaccard-Tanimoto (JT),^14,15^ Tversky’s,^16^ to mention few. In the cheminformatics community, the Jaccard-Tanimoto (or simply Tanimoto) is the preponderant similarity index.^17^ Equations 1, 2, and 3 show the formulas for the Jaccard-Tanimoto, Russell-Rao, and Sokal-Michener indexes; respectively. Where *a* represents the one-one coincidences, *b* the zero-one mismatches, *c* the one-zero mismatches, and *d* the zero-zero coincidences.^1^

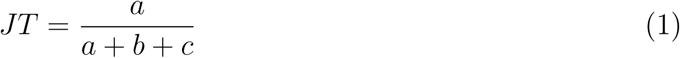

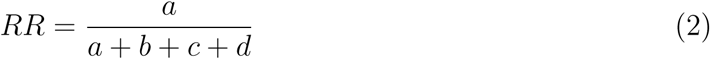

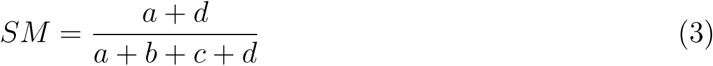

The traditional way of calculating similarity is by comparing two molecules. In this way, to calculate the average similarity of a dataset with *N* molecules, the *N* ^2^ matrix of comparisons would be needed.^18^ The calculation of complete similarity matrices is used in the quantification of diversity^18^ and in clustering algorithms like Taylor-Butina. ^19^ Recently, the iSIM (instant similarity) framework was introduced as a way of quantifying the average pairwise similarity of a dataset in linear scaling complexity *O*(*N* ).^20^ The iSIM framework simply adds the fingerprints resulting a in vector [*k*_1_*, k*_2_*, …, k_i_*], where each *k_i_* will represent the number of ones in each column. It is easy to get the number of one-one coincidences per column, 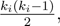 , after adding for all *i*s, we obtain a value that we can substitute for the *a* in the similarity indexes of choice. Analogously, expressions for *b*, *c*, and *d* can also be easily obtained.^20^ For example, the RR index could be transformed to equation 4, with an expression we can obtain the exact same value as the average of the pairwise operations, without having to compute them.

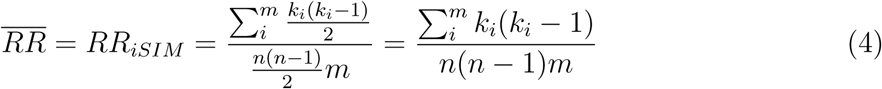

Even though getting the iSIM value for a data set of compounds is extremely useful in several tasks like diversity selection, chemical space exploration,^20^ and clustering;^21^ it is not enough information to describe the whole distribution of all the pairwise similarities in a set, reason that impulsed us to develop iSIM-sigma.

As the number of molecules in chemical libraries increases, the pairwise approach of calculating the variance of the similarities is unfeasible. In this work, we show a fast alternative way of approximating the standard deviation of the pairwise molecular similarities without the need to compute them. The proposed method uses a stratified sampling strategy based on complementary similarity. We also show that is possible to obtain the exact value for indices like RR and SM in a less computationally costly way for large libraries. Intending to show the relevance of knowing the standard deviation, we show an example where we calculate the standard deviations of selected molecules through different selection methods and how they compare to the whole dataset. On this line, we also introduce iSIM-sigma as a measurement of cluster compactness, we compare it with other used metrics for this task.

## Theory

### Exact approach for RR and SM

As the reader can notice, the iSIM RR calculation corresponds to the probability of a one-one bit coincidence.^20^ The simplest way of approximating the standard deviation of the comparisons would be to treat the comparisons as a simple Bernoulli distribution. ^22,23^ Since we already have the probability of a successful event (*RR_iSIM_* ), we could estimate the standard deviation using the normal approximation for a population sample proportion,^24^ as shown in equation 5. Where our sample size would correspond to the length of the fingerprints, *m*, since we follow a column-wise approach. When we try this approach on randomly generated fingerprints the approximation is quite accurate (consult SI). However, since real-molecule fingerprints are not randomly encoded, this approach fails to approximate the standard deviations of the comparisons (consult SI).

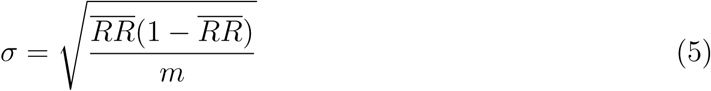

It is clear that more information about the distribution of ones across the columns is needed. Fortunately, by considering pairs of columns, we can obtain all the information necessary to compute the standard deviation.

For a given pair of fingerprints, *a* is defined according to the following formula.

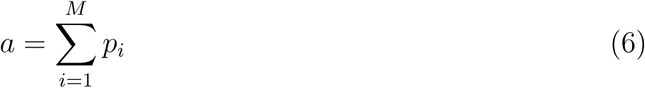

Where *p_i_*is 1 if both molecules have a value of 1 at position *i*, and 0 otherwise. For iSIM, to compute the average value of *a*, *E*(*a*), we used linearity of expectation according to equation 7, where *E*(*p_i_*) is simply [inline] .

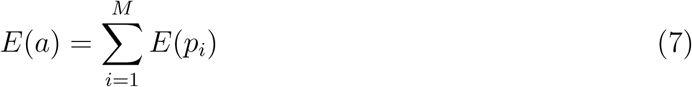

The variance of *a* doesn’t follow this same linearity but instead expands to include co- variances between terms according to equation 8, where *σ_pi,pj_* is the covariance between *p_i_*and *p_j_*.

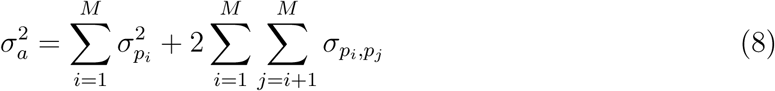

Since *p_i_*can only have a value of 0 or 1, it follows a Bernoulli distribution and has a variance following equation 9.

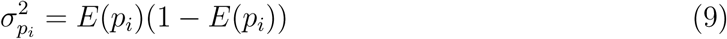

We additionally note that covariance is defined according to equation 10.

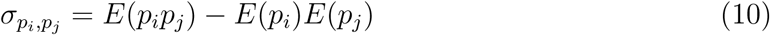

From equations 8, 9, and 10, we can therefore compute *σ*^2^ if we know *E*(*p_i_p_j_*) for any pair of *i* and *j*, and *E*(*p_i_*) for any *i*. *E*(*p_i_*), as previously mentioned, is computed from iSIM and takes *O*(*N* ) time. *E*(*p_i_p_j_*) is simply the probability that when selecting a pair of two molecules, there is a value of 1 for both molecules at positions *i* and *j*. Thus, similar to how we computed *E*(*p_i_*), we go through each of the molecules and enumerate the number of instances where there is a 1 in both positions *i* and *j*, defined as *α_ij_*. Since there are 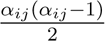 pairs where *p*_1_*p*_2_ is 1, and 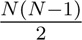 total pairs, *E*(*p p* ) simply follows equation 11.

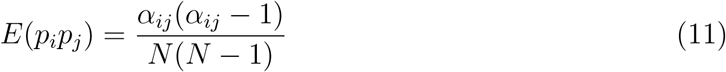

Determining *α_ij_* takes *O*(*N* ) time, and we must do this for all *O*(*M* ^2^) pairs, thus, determining all the covariances takes *O*(*NM* ^2^) time. We can therefore obtain the variance of *a* in *O*(*NM* ^2^) time, and all we need to do is square root this, and then divide it by *m* to get the standard deviation of the RR metric.

For the SM metric, we take a similar approach, with

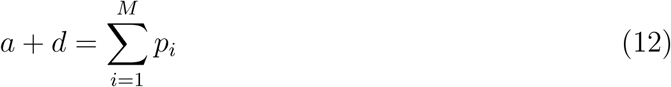

Where *p_i_*is now 1 if both molecules have a value of 0, or both molecules have a value of 1 for the fingerprint, and 0 otherwise.

To compute *E*(*p_i_p_j_*), we now need to consider four separate cases: both molecules have a value of 1 for positions *i* and *j*, both molecules have a value of 0 for positions *i* and *j*, both molecules have a value of 1 for position *i*, and 0 for position *j*, and both molecules have a value of 0 for position *i*, and 1 for position *j*. Similar to the RR metric, we go through each of the fingerprints and enumerate the number of instances where there is a 1 in both positions *i* and *j* (*α_ij_*), the number of instances where there is a 1 in position *i* and a 0 in position *j* (*β_ij_*), the number of instances where there is a 0 in position *i* and a 1 in position *j* (*γ_ij_*), and the number of instances where there is a 0 in both positions *i* and *j* (*δ_ij_*). *E*(*p_i_p_j_*) then follows equation 13. Determining *α_ij_*, *β_ij_*, *γ_ij_*, and *δ_ij_* for all pairs of *i* and *j* will take *O*(*NM* ^2^) time.

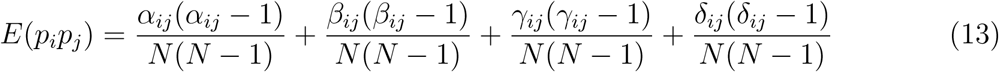

We can use this to obtain the variance of *a* + *d*, and then take its square root and divide by *m* to get the standard deviation of the SM metric.

This approach is able to get the exact solution for the calculation of the standard deviation in a *O*(*NM* ^2^) scaling, which corresponds to a huge improvement for ultra-large libraries where *O*(*N* ^2^) *>> O*(*NM* ^2^). In cases where the denominator of the similarity index is constant for all the comparisons (eg. RR or SM), the same reasoning can be applied to get the same result. Those indexes are going to be easier to approximate since only the variances and covariances of the numerator are needed. For the JT index, this approach does not obtain good results. The denominator is not constant across all the comparisons, there are also variances and covariances needed from the denominator. In literature, modeling of JT similarities distribution has been done but for comparing one query to the rest of the set in the light of similarity searches significance.^22,23^ In those approaches variances and covariances for the denominator and numerator are modeled, but the calculation is not exact^23^ and is not for all the pairwise similarities. For indexes, like JT, where the exact approach does not work, in the next section, we present a sampling method to estimate their standard deviation.

### Stratified sampling approximation

One approach to estimate the standard deviation of a distribution of similarity scores is by sampling representative molecules that can unveil the whole distribution properties. The most common approach is to sample random molecules ^25^ from the set, this method has the disadvantage of not being deterministic, hence some random sampling may yield a bad sampling. Other popular selection methods are MaxMin,^26^ MaxSum;^27^ for those, the complete similarity matrix is needed; then there would be no point in using them to estimate the standard deviation. Recently, with our iSIM framework, we included several selection methods based on complementary similarity.^20^ The complementary similarity is the similarity of the remaining set when one molecule is removed from it. This value is going to tell how similar was the removed molecule to the rest of the set, therefore it can be used to rank the molecules from the medoid (the most similar to the rest of the molecules, low complementary similarity) to the outlier (the least similar to the rest of the molecules, high complementary similarity). Since, iSIM is able to calculate the similarity of a whole set linearly, calculating the complementary similarity for each molecule in the set will be also an *O*(*N* ) step. Figure 1 illustrates the complementary similarity ranking procedure for a small set of molecules.

**Figure 1:**
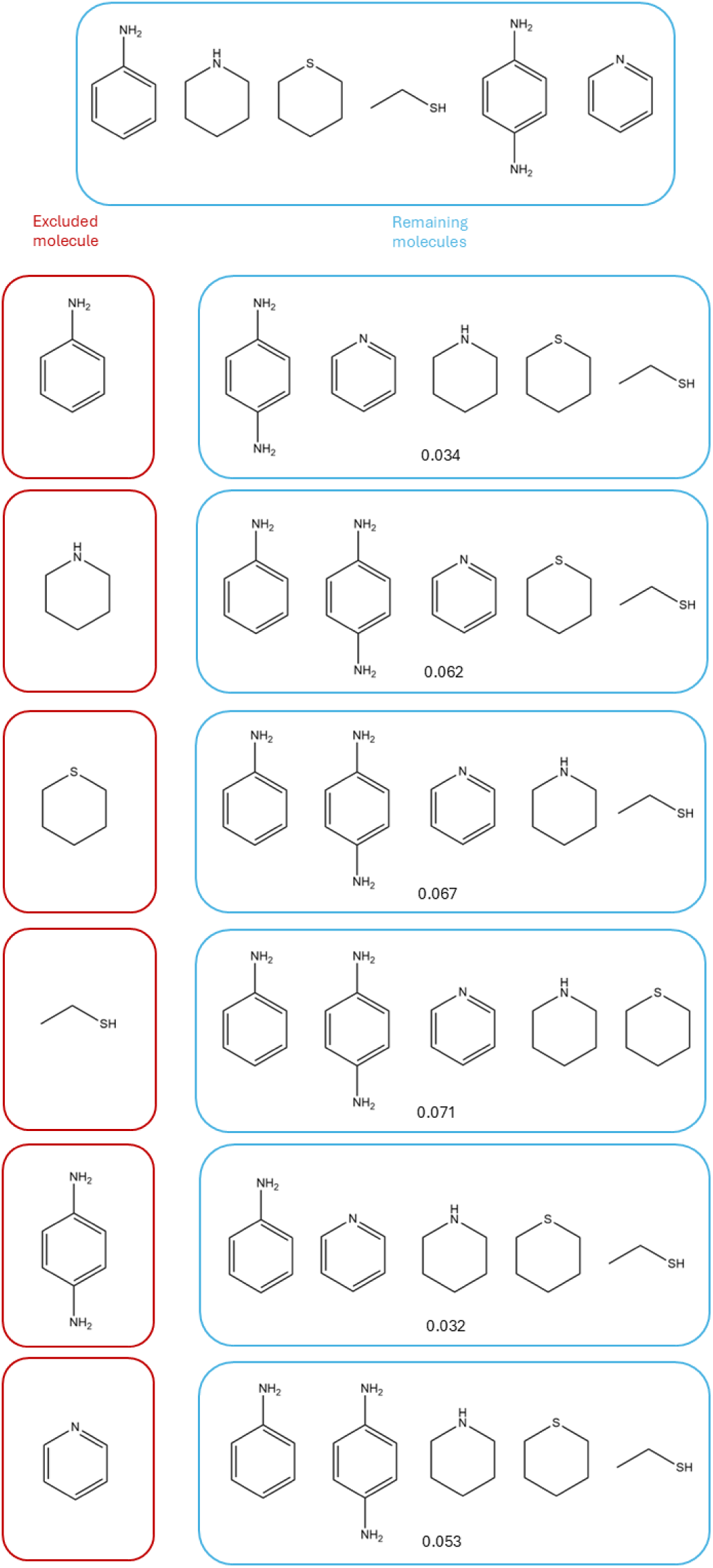
Example of iSIM complementary similarity ranking procedure. A molecule is excluded from the dataset, the iSIM (average similarity) of the remaining molecules is calculated. That value corresponds to the complementary similarity of the removed molecule. Molecules were represented using ECFP6 fingerprints and compared with iSIM Tanimoto.

The complementary similarity ranking can be used to select a representative set. We previously introduced stratified sampling, which divides the complementary similarity ranking into a desired number of bins, *b*, all with the same number of molecules. Then a fixed number of molecules are picked from each bin (depending on the number of molecules the user wants to sample). The sampled molecules will cover all the complementary similarity range. It was showed how the average similarity of the sampled molecules was close to the average similarity of the whole set, and also how their first two principal components resemble the space of the whole set. ^20^ For those reasons, we chose stratified sampling as a possible way of estimating the standard deviation of the similarity distribution of the whole set. In Figure 2 we continue with the example from Figure 1 and show graphically how stratified sampling works.

**Figure 2:**
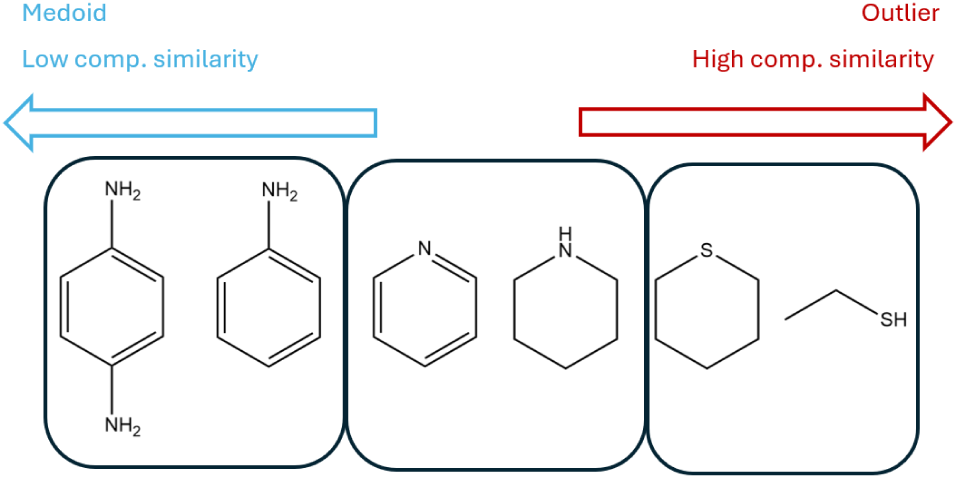
Stratified sampling example, using the molecules from Figure 1. The molecules are sorted in ascending complementary similarity. For three molecules to sample, the molecules are separated in bins and one molecule per bin is sampled.

The stratified sampling iSIM-sigma approach starts by sampling an *n* number of molecules from a dataset of size *N* . Then the pairwise comparisons of the *n* (10, 25, and 50 in this work) molecules are calculated and subsequently the standard deviation of those comparisons.

## Results and discussion

### Exact approach for RR

We tested our *O*(*NM* ^2^) exact method on 500 subsets of the ChEMBL33 library with random sizes ranging from 1000 to 5000 molecules. Figure 3 shows the comparison of our method with the pairwise standard deviation calculation for three similarity indexes. As it can be seen the fit is perfect for the *RR* and *SM* indexes; but not for the *JT* index. For the former two indexes, in their definition (equations 2 and 3) their denominators are constant across all comparisons since the number of bits in all the molecules in the set is the same. This allows us to only care about the numerator’s distribution. As shown in the theory section, it is possible to obtain the standard deviation of the pairwise comparisons, with only the variances and covariances of the columns. The perfect fit for those two indexes supports our mathematical formulation. Despite the exact calculation of the standard deviation of the similarities for those indexes, this formulation does not work for indexes like *JT* , since the denominator in the comparisons is not always the same (equation 1).

**Figure 3:**
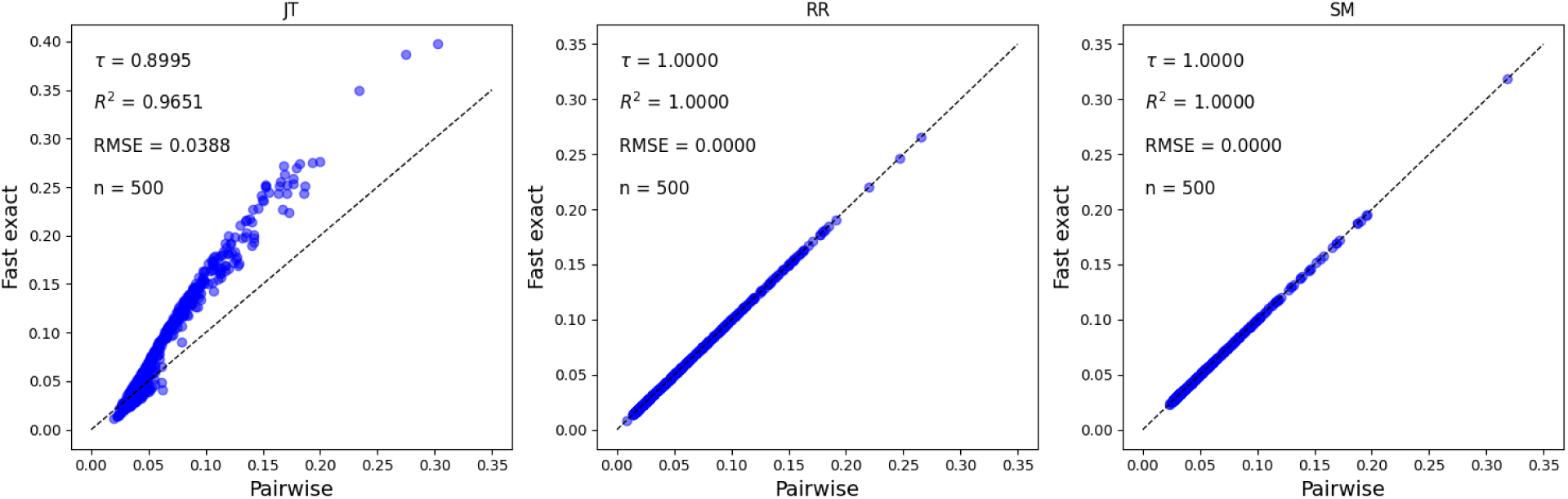
Comparison of the pairwise standard deviation of the similarity values and the fast exact approach for 500 randomly selected subsets (sizes between 1,000-5,000) of the ChEMBL33 library represented with RDKit fingerprints (2048 bits); using Jaccard-Tanimoto (JT), Russell-Rao (RR), and Sokal-Michener (SM) similarity indexes.

To resolve this issue, we turned our way into a sampling strategy to estimate the standard deviation. We used the above-explained stratified sampling method to select representative molecules, then calculate the pairwise comparisons between the selected molecules and calculate their standard deviation. To show the importance of selecting representative molecules, we calculated the same approach but selected the molecules randomly. Figure 4 shows the results for the sampling approaches using the *JT* index. When comparing the three stratified methods, is clear that the higher the number of selected molecules the standard deviation estimation is going to be closer to the actual. We can see that when selecting 50 representative molecules, the *RMSE*, *Kendall* − *Tau*, and *R*^2^ indicate that the approach is overall solid. We can see that the random approach, despite also selecting 50 molecules, has several deviations from the actual values, and they actually have worse metrics than when selecting only 25 molecules with the stratified approach. With this, we can infer that selecting representative molecules is important to estimate the standard deviation of the similarities of the whole set. Random methods could perform well, but it ends up being a chances game, they are unstable.

**Figure 4:**
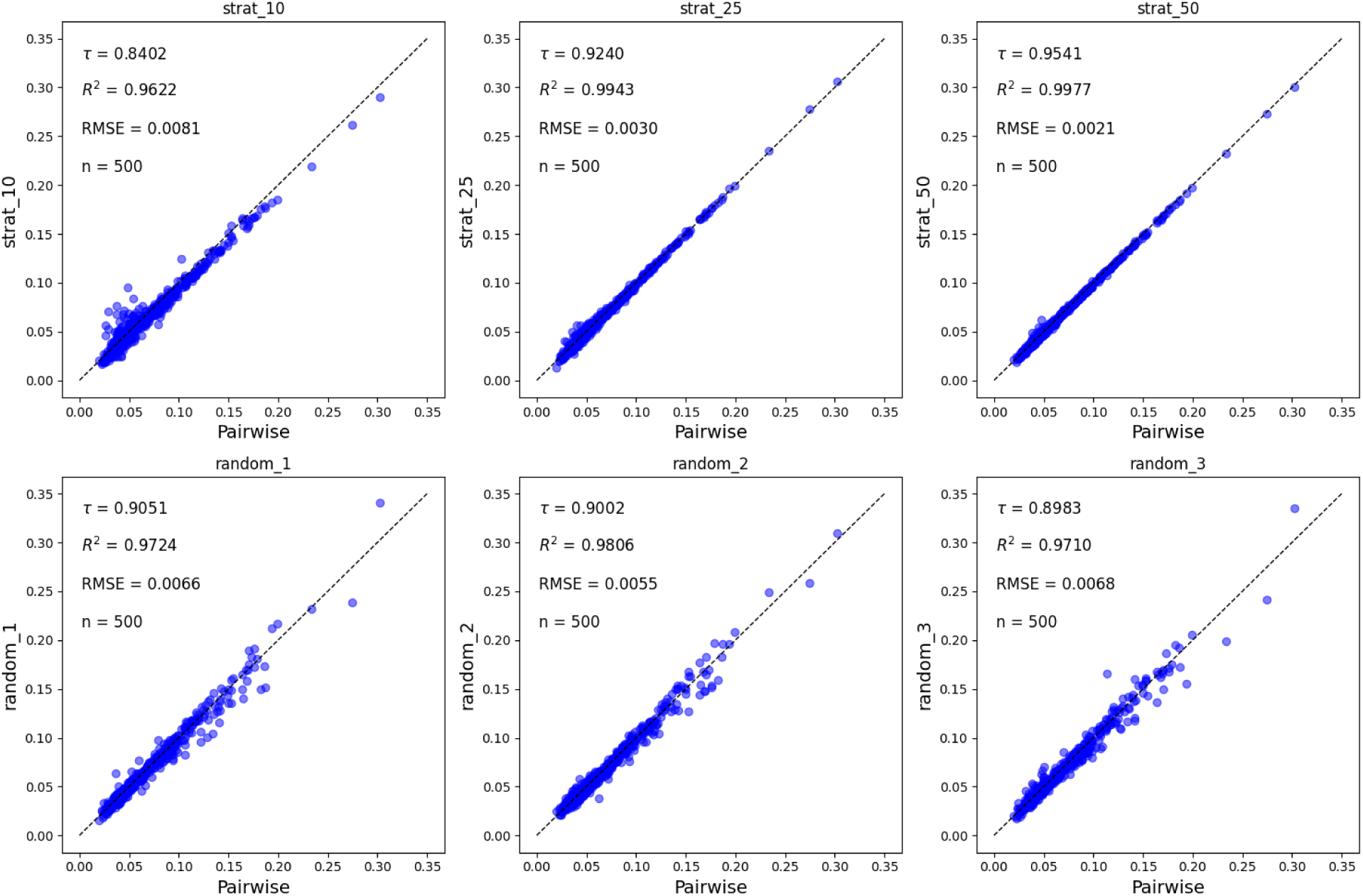
Comparison of the pairwise standard deviation of the similarity values and sampling approaches for 500 randomly selected subsets (sizes between 1,000-5,000) of the ChEMBL33 library represented with RDKit fingerprints (2048 bits) using the Jaccard-Tanimoto (JT) index. Stratified methods use 10, 25, and 50 sampled molecules respectively. Three trials of a random selection of 50 molecules are shown.

**Figure 5:**
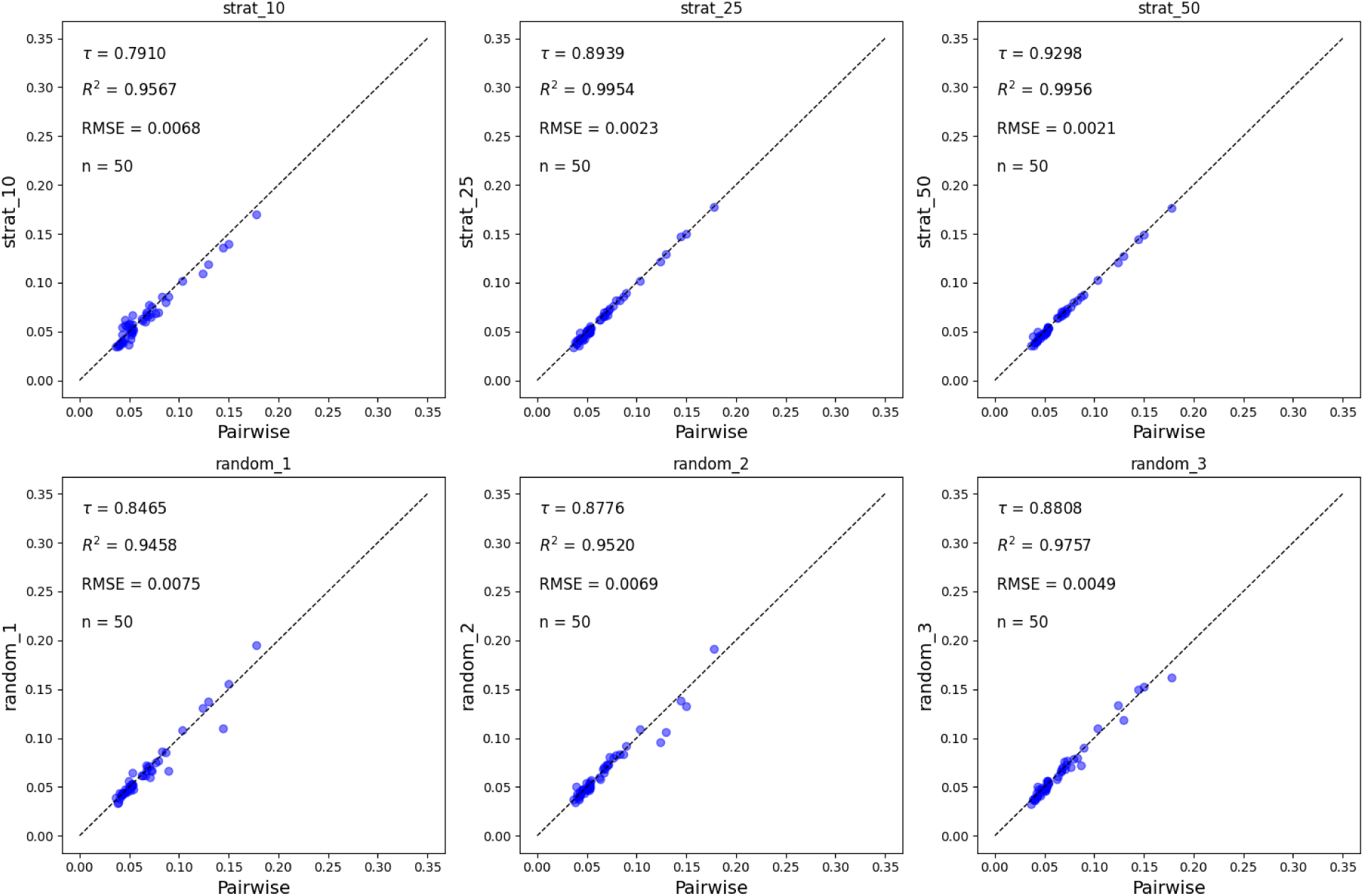
Comparison of the pairwise standard deviation of the similarity values and sampling approaches for ChEMBL33 subsets, with sizes between 1,000 and 50,000, represented with RDKit fingerprints (2048 bits) using the Jaccard-Tanimoto (JT) index. Stratified methods use 10, 25, and 50 sampled molecules respectively. Three trials of a random selection of 50 molecules are shown.

With the aim of studying how the sampling methods behaved on bigger datasets, we tested them on ChEMBL33 subsets increasing the size in steps of 1000 molecules, getting up to 50,000. On 5 we show the results for these 50 sets, it is clear that the strong performance of the stratified method sampling 50 molecules remains. While the random sampling methods underperform in all three metrics. Figure 6 shows the relation between the size of the subset and the absolute errors. It can be appreciated how by increasing the number of sampled molecules the absolute errors get lower, in fact for the stratified method when sampling 50 molecules only two of the subsets have errors higher than 0.005, but all errors are lower than 0.01. This is impressive and solidifies stratified sampling as a good alternative to select representative molecules from a set. With only 50 molecules, we can obtain the variance in similarities of a whole data set. We can also notice how the random sampling method varies from trial to trial, and for some sets, the errors are higher than 0.01, and a considerable number are above 0.005. This solidifies how random sampling will not be always a reliable way of estimating the standard deviation of the similarities.

**Figure 6:**
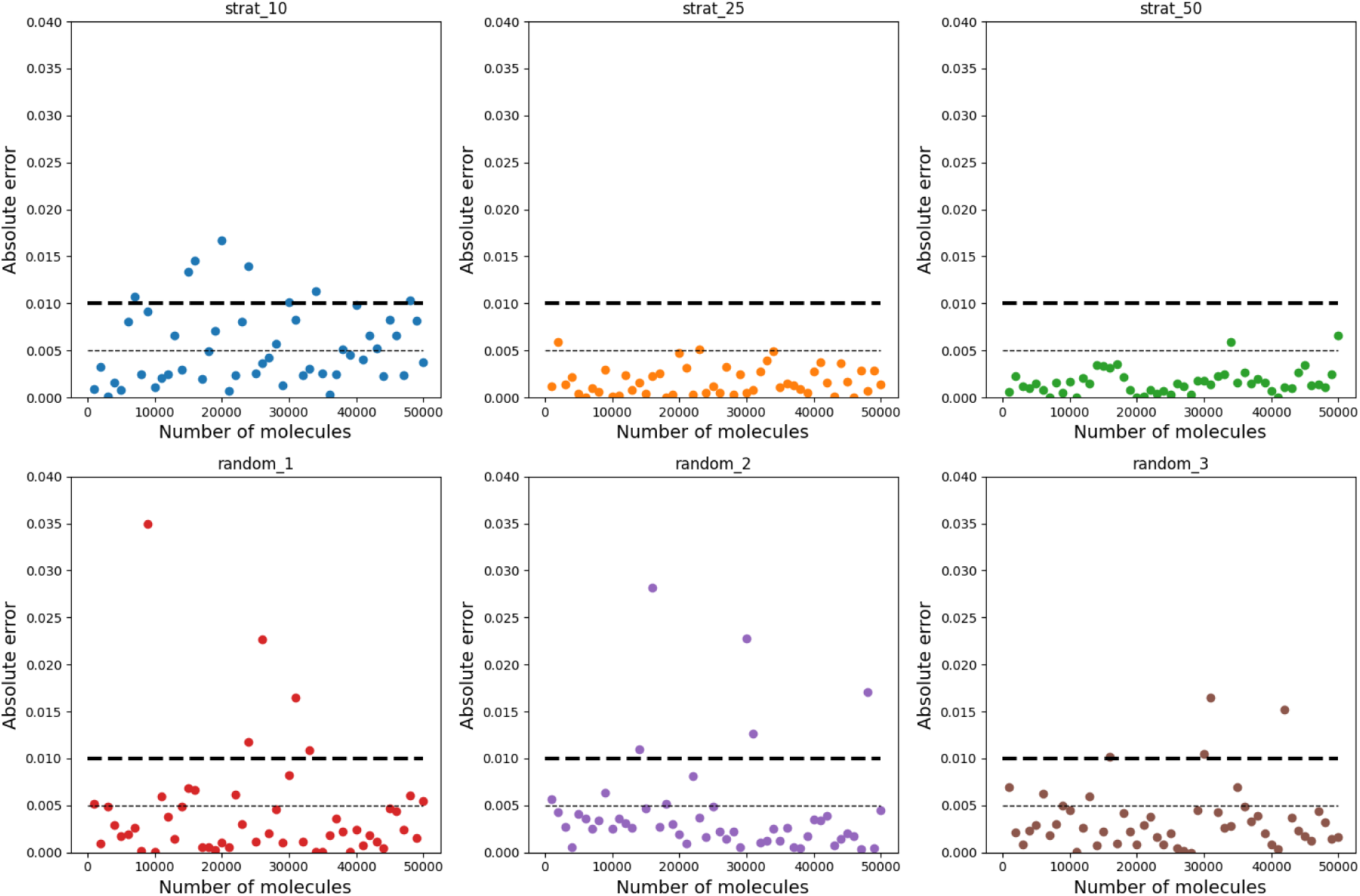
Relationship between the absolute errors and size of data set for the stratified and random sampling standard deviation estimation methods for subsets of ChEMBL33 library represented by RDKit fingerprints (2048 bits). Sets vary in size by 1,000 molecules, from 1,000 to 50,000 molecules.

**Figure 7:**
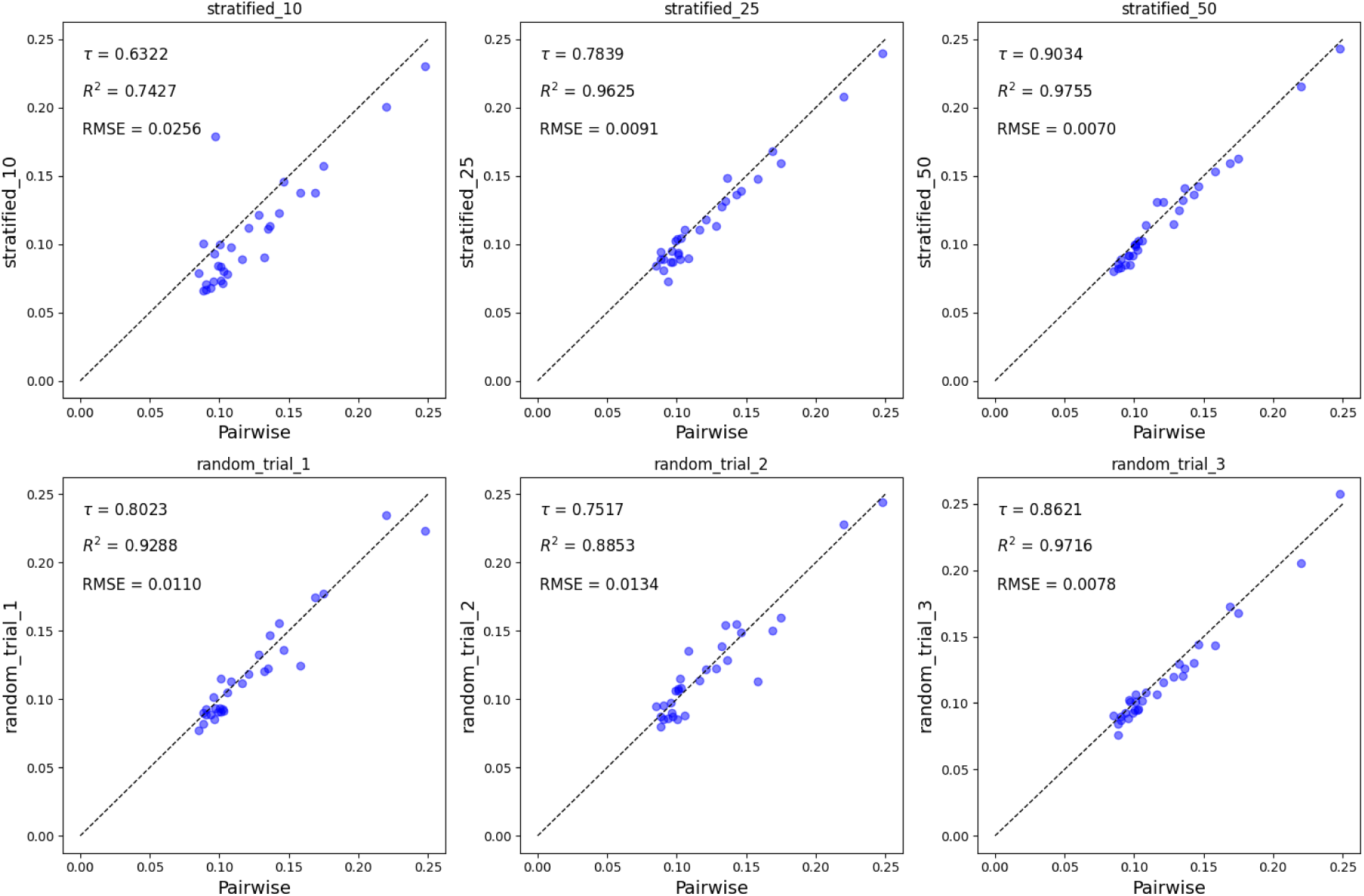
Comparison of the pairwise standard deviation of the similarity values and sam- pling approaches for 30 data sets each with compounds with activity against a different biological target using the Jaccard-Tanimoto (JT) index. Molecules represented with RDKit fingerprints (2048 bits). Stratified methods use 10, 25, and 50 sampled molecules respectively. Three trials of a random selection of 50 molecules are shown.

**Figure 8:**
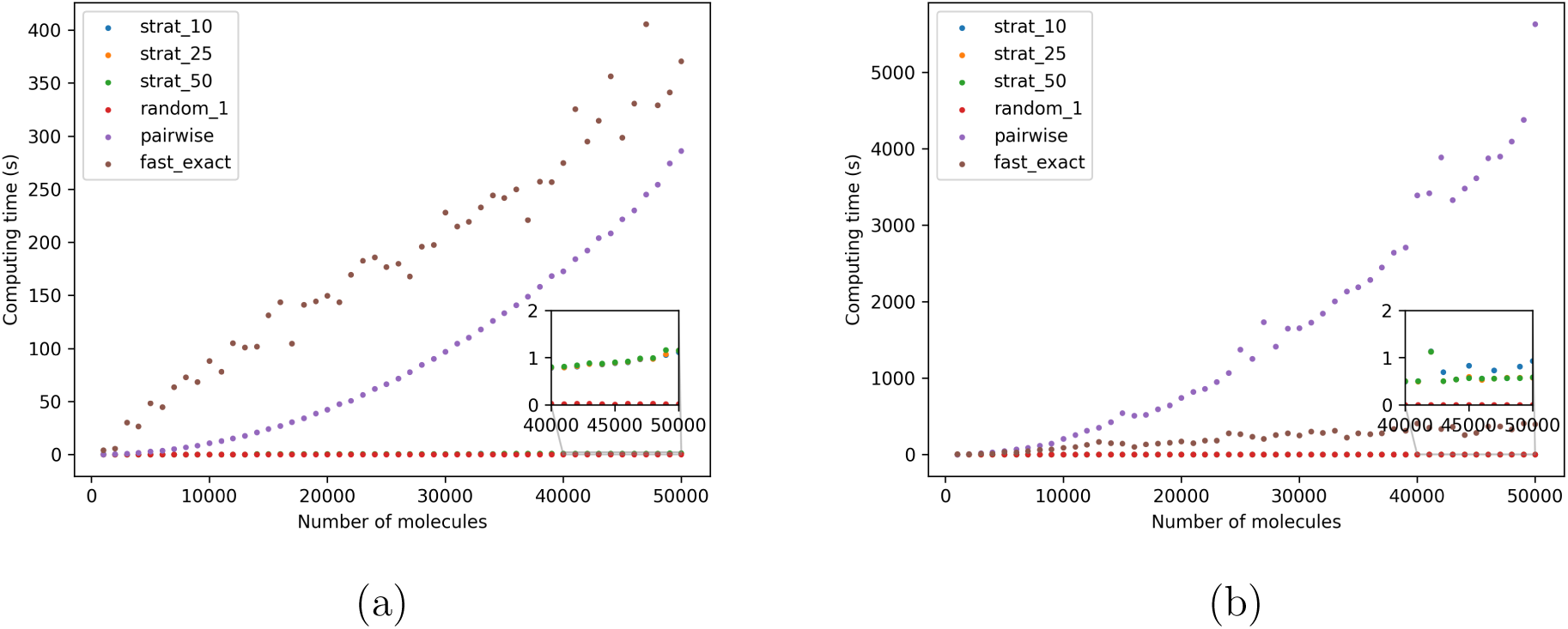
Computing time versus number of molecules in the dataset for each standard deviation estimation method using (a) *JT* and (b) *RR*. Both are calculated for sets of the ChEMBL33 library represented with RDKit fingerprints (2048 bits).

**Figure 9:**
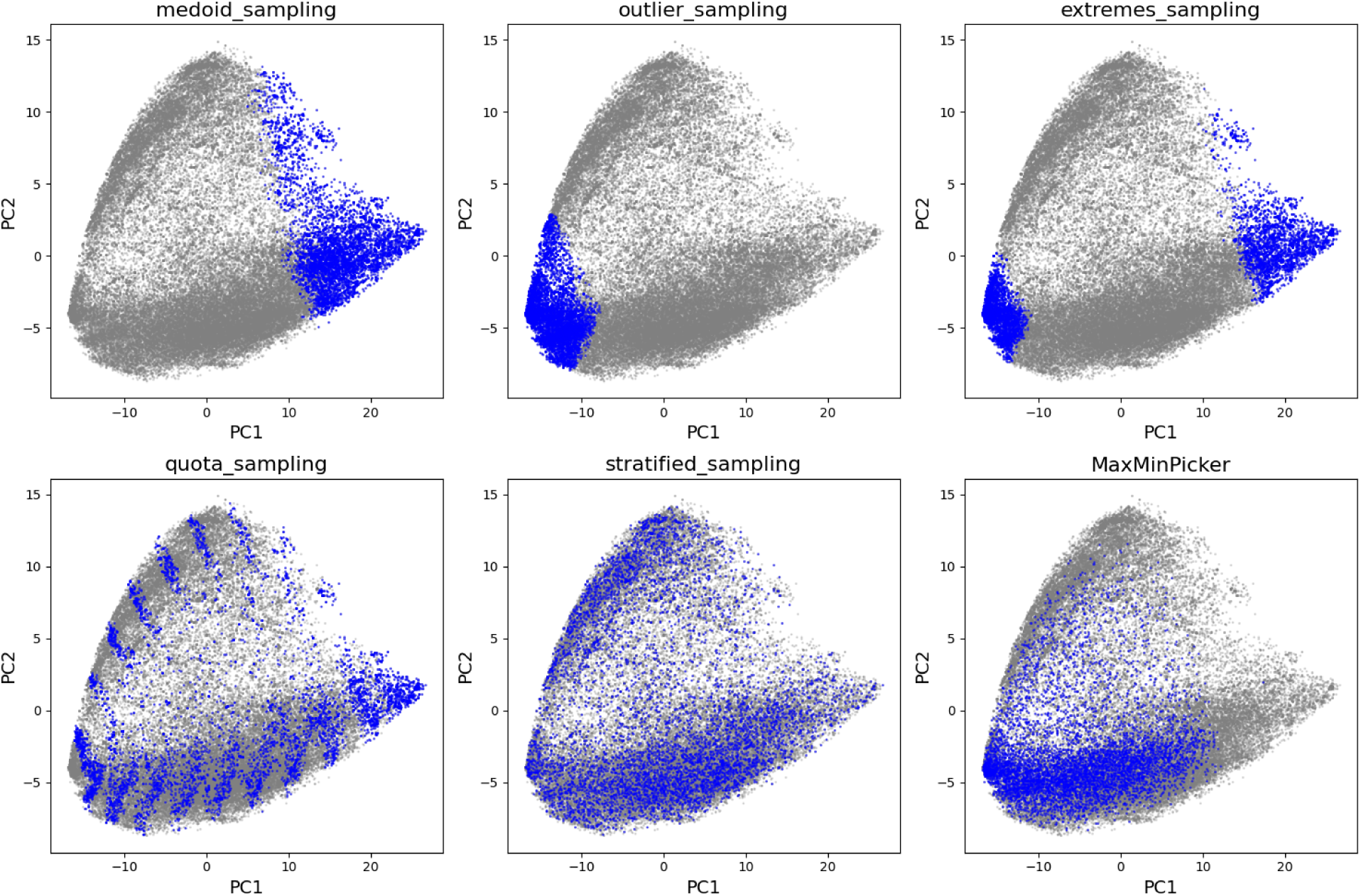
Scoring plots for the first two principal components of ChEMBL33 natural products set (n = 64086), represented with RDKit fingerprints (2048 bits). Selected molecules are colored in blue, and non-selected molecules in gray.

So far we have shown results from subsets of the ChEMBL library, but we also tested our methods on higher structured data. For 30 curated sets of molecules with activity for different biological targets, we evaluated the performance of the proposed sampling methods. We demonstrate again how the stratified sampling method with 50 sampled molecules is the one that can estimate standard deviations with lower errors. Is fair to mention that one of our random trials has a performance close to our stratified method, however, the other two trials have higher errors; confirming the unreliability of random sampling from trial to trial. These trends across the methods are followed for other types of fingerprints, in our supporting information we show how for ECFP4 and MACCS fingerprints iSIM-sigma can still estimate the standard deviation with low errors. iSIM-sigma holds its accuracy with other similarity indexes like *RR* and *SM* , as shown in our supporting information. In smaller datasets or shorter fingerprints, it would make sense to use the exact approach for these two indexes.

We continue our discussion by comparing the timing of all the shown methods. Figure 10 (a) shows the computing times for each method when estimating the standard deviation of *JT* similarities of the whole set. As expected the pairwise approach takes a quadratic form, while the sampling methods are incredibly faster than. Even though the random sampling method is the fastest and constant, we have seen that its accuracy is not always good. The sampling methods can accurately estimate the standard deviation for 50,000 molecules in about one second, which seems like a fair time to pay considering its superior accuracy to the random sampling. We want to point out that the pairwise method for the *JT* is faster than the exact approach because we used the C++-optimized bulk similarity function from RDKit to calculate the pairwise similarities. When comparing simple numpy functions, like the ones we use for our sampling methods, we can see that the fast approach is faster than the quadratic scaling pairwise way, we can observe this in the timing for the *RR* in Figure 10 (b).

**Figure 10:**
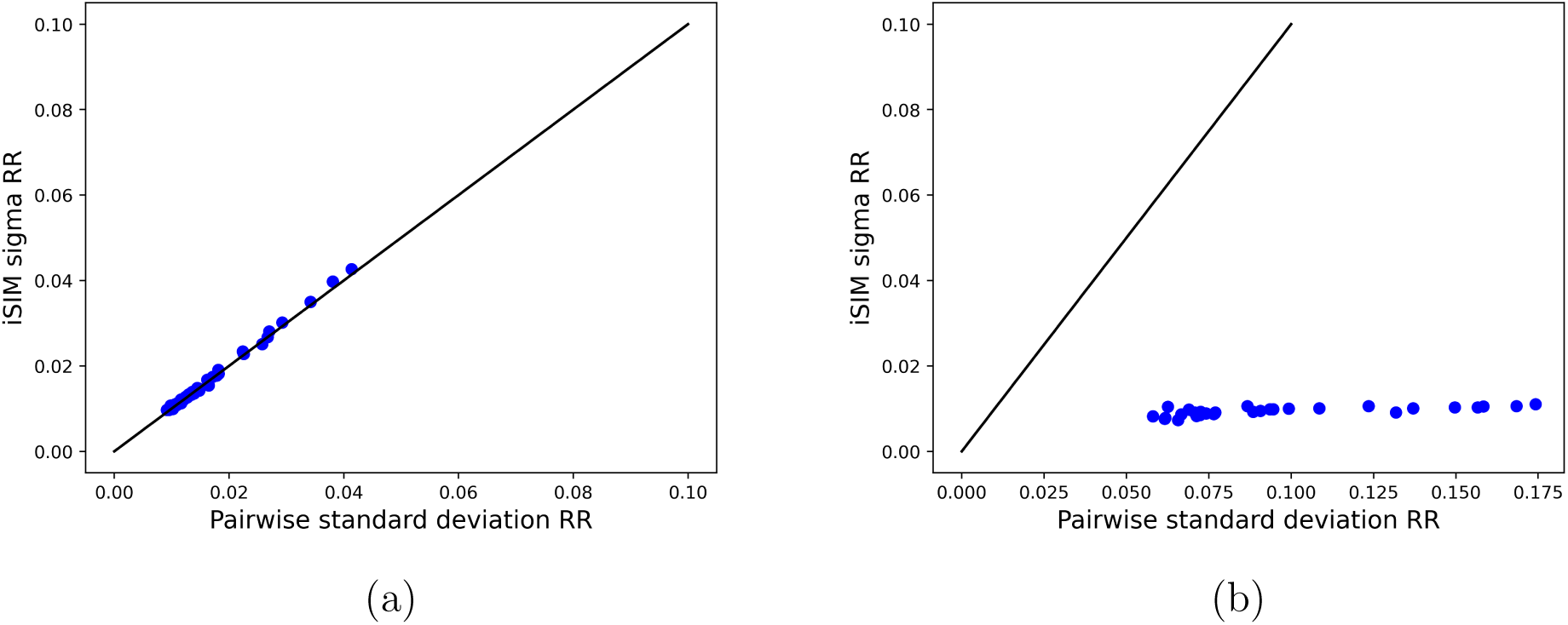
Pairwise standard deviation versus Bernoulli binomial normal approximation approach for (a) randomly generated fingerprints and (b) ChEMBL target-specific libraries represented with RDKit fingerprints.

Having shown the high accuracy of the proposed methods, now we want to exemplify one application of the fast estimation of the standard deviation. iSIM-sigma can be used to estimate the standard deviation of sampled molecules with different sampling methods. The user can compare the values of the average similarity (iSIM) and the standard deviation of similarities (iSIM-sigma) for the selected molecules to the whole set’s values. In this way, it can be assessed if the sampled molecules’ chemical space resembles the whole dataset. We tested our approach on the ChEMBL33 natural products dataset, we tested six different sampling algorithms (5 based on iSIM complementary similarity), and RDKit’s fast implementation of MaxMin. In Table 1, we show the iSIM, iSIM *σ*, and pairwise *σ* for the selected 10% with each method. As it can be appreciated, the *σ* estimation is pretty accurate for most, it only fails to estimate for the outlier methods (error higher than 0.01). With these values, we can see that stratified sampling is the only one that reproduces the distribution variables when compared to the values of the whole set. The values for MaxMin reinforce that the algorithm’s goal is to minimize the similarity of the selected molecules, but not having close average and variances to the whole dataset; since their values are far from close to the whole set’s.

**Table 1:** iSIM JT, iSIM-JT, and Pairwise *σ* for sampled molecules (10%) from ChEMBL33 natural products (whole dataset: n = 64086, iSIM = 0.2745, iSIM-*σ* = 0.1302) using different sampling methods. Molecules were represented with RDKit fingerprints (2048 bits).

| Sampling method | iSIM | iSIM $\sigma$ | Pairwise $\sigma$ |
| --- | --- | --- | --- |
| Medoid | 0.6031 | 0.0747 | 0.0763 |
| Outlier | 0.0856 | 0.0687 | 0.0807 |
| Extremes | 0.2800 | 0.2742 | 0.2748 |
| Quota | 0.3113 | 0.1913 | 0.1890 |
| Stratified | 0.2745 | 0.1329 | 0.1302 |
| MaxMin | 0.1764 | 0.0936 | 0.0900 |

Visually we can see in Figure 9 how the different methods sample molecules and observations are consequent with the obtained values for their *σ*. Medoid and Outlier samples contain a compact Chemical Space, which agrees with them having the lowest iSIM-sigma standard deviations. Extremes sampling samples from both medoids and outliers regions, making the distribution of similarities have a high variance. The extremes sampling is one situation where is key to know the variance of the distribution, as the average similarity of the sampled molecules is close to the whole set’s, but clearly, the standard deviation indicates that the sampled molecules do not resemble the entire chemical space. The iSIM and iSIM-sigma calculations can be useful in instances of diversity selection, or data splitting.

## Methods

All scripts used for this work are included in the repo https://github.com/mqcomplab/ iSIM_sigma.git. Functions to calculate the faster exact approach and the stratified iSIM-sigma are included in the isim_sigma.py script. Main calculations were done using RDKit^28^ fingerprints (2048 features), and results for MACCS^3^ and ECFP4^8^ are included in the SI. For the subsets calculations ChEMBL33 library ^29^ was used, the subsets were generated by selecting two random sections of the database with the molecules ordered by population count and combining them. This approach was taken because a simple random selection gives virtually the same variance in each subset. Subsets were selected in increasing sizes from 1,000 to 50,000 fingerprints. The target-specific libraries were obtained from van Tilborg et al.^30^ For the sampling example, the natural products of the ChEMBL33 library were used, and compressed fingerprints are included in the repository. Jupyter notebooks with examples of how to obtain the shown results are included in the repository.

## Conclusions

In this work, we showed how it is possible to calculate the exact standard deviation of the similarity distribution for similarity indexes like RR and SM without the computation of the pairwise similarity matrix. Even though this approach is a huge improvement in the complexity of the algorithm, it has two main drawbacks: it is not applicable to all similarity indexes and would not be as efficient for high-dimensional representations. To overcome those problems we propose a sampling-based estimation method for the standard deviation of the similarities, which we call iSIM-sigma. By selection 50 representative molecules were are able to estimate the standard deviation with RMSEs below 0.01 units for randomly selected subsets of up to 50,000 molecules from the ChEMBL library, and also for target-specific sets. The stratified approach has linear complexity and shows lower errors than the random sampling alternative. We also showed that iSIM-sigma can be useful for calculating the variances of similarities in selected subsets, and how it gives extra key information.

## Acknowledgement

KLP, BZ, and RAMQ thank the National Institute of General Medical Sciences of the National Institutes of Health for support under award number R35GM150620.

## Supporting Information Available

**Figure 11:**
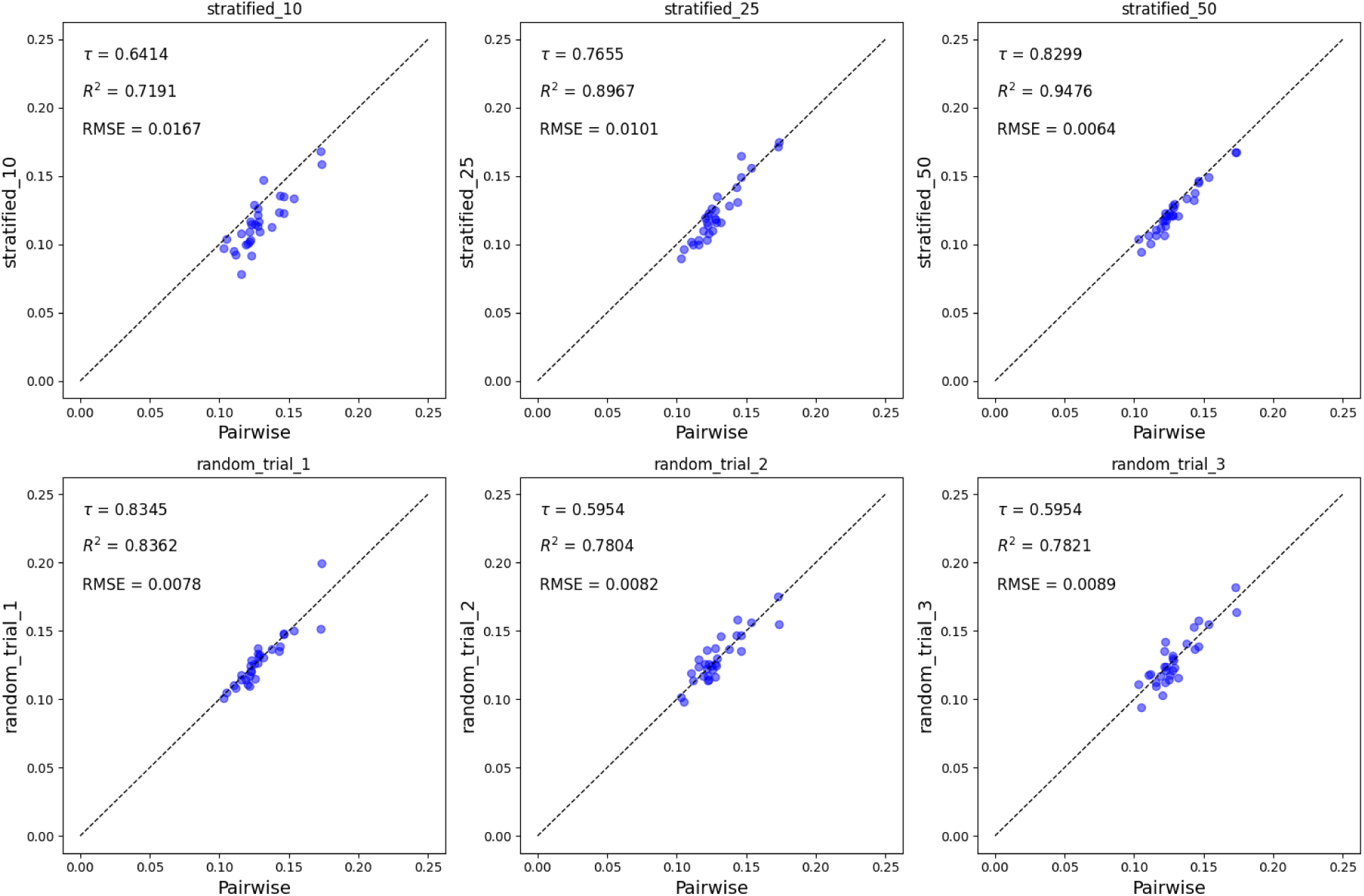
Comparison of the pairwise standard deviation of the similarity values and sampling approaches for 30 data sets each with compounds with activity against a different biological target using the Jaccard-Tanimoto (JT) index. Molecules represented with MACCS fingerprints (167 bits). Stratified methods use 10, 25, and 50 sampled molecules respectively. Three trials of a random selection of 50 molecules are shown.

**Figure 12:**
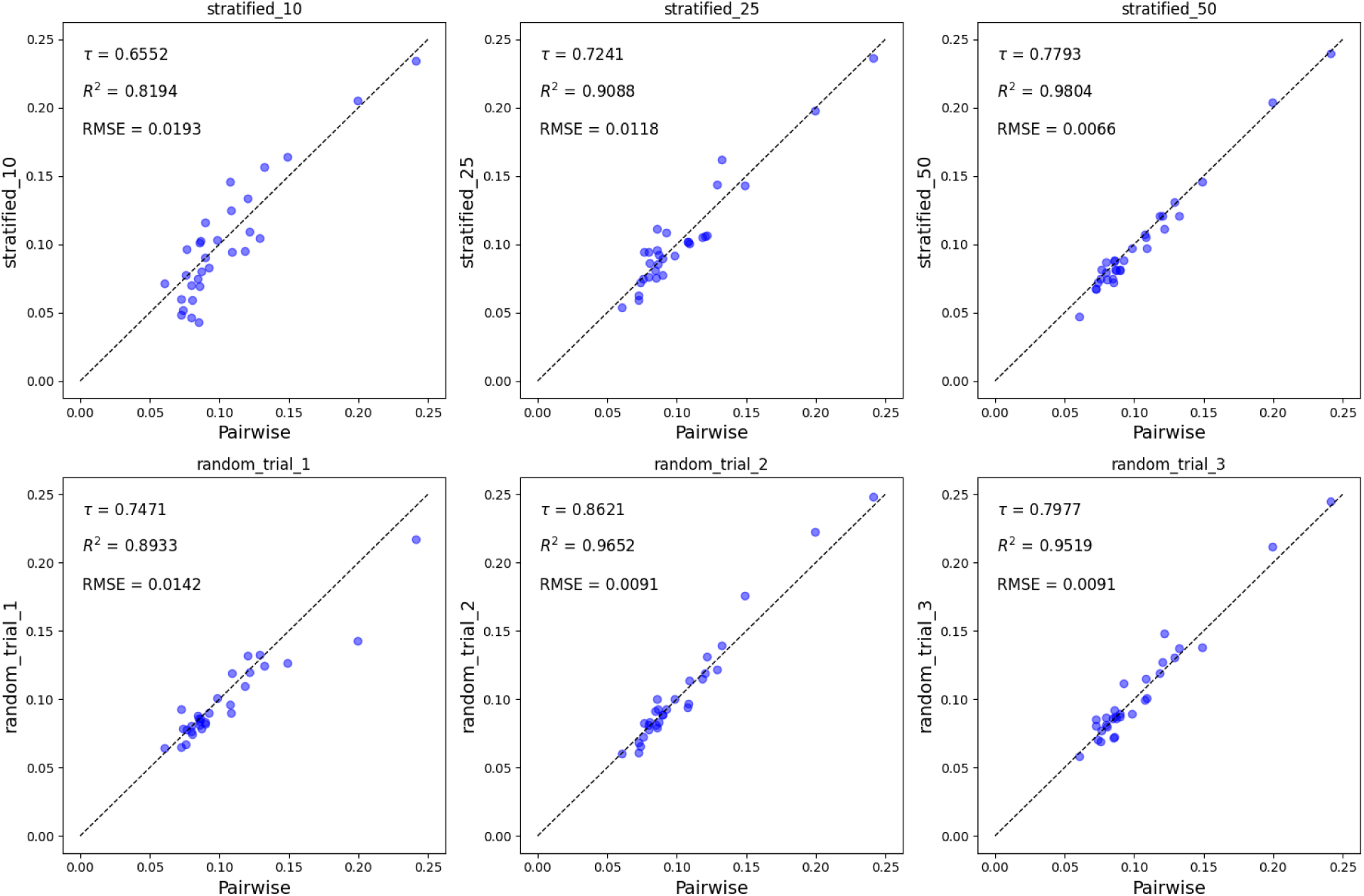
Comparison of the pairwise standard deviation of the similarity values and sampling approaches for 30 data sets each with compounds with activity against a different biological target using the Jaccard-Tanimoto (JT) index. Molecules represented with ECFP4 fingerprints (1024 bits). Stratified methods use 10, 25, and 50 sampled molecules respectively. Three trials of a random selection of 50 molecules are shown.

**Figure 13:**
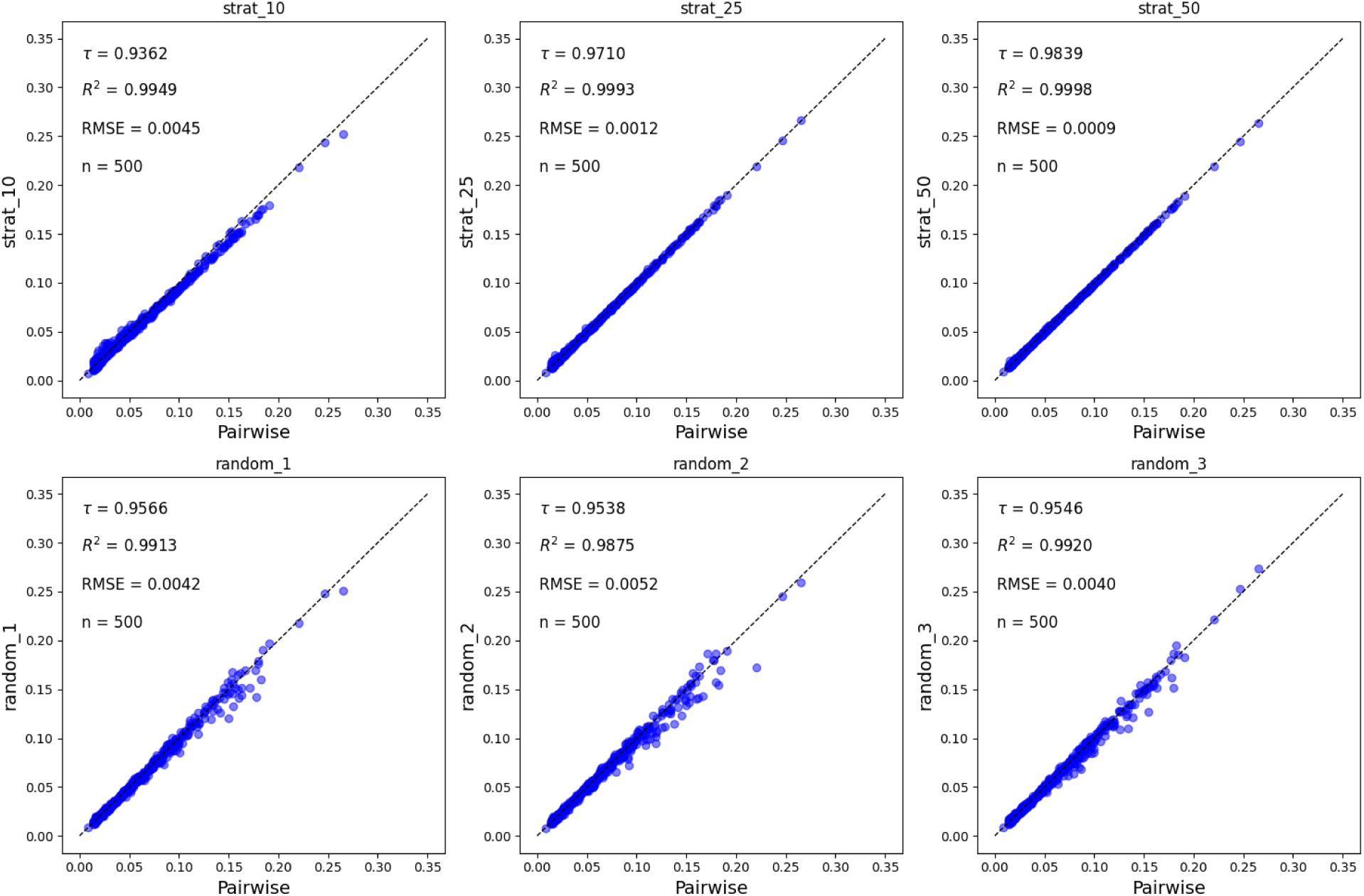
Comparison of the pairwise standard deviation of the similarity values and sampling approaches for 500 randomly selected subsets (sizes between 1000-5000) of the ChEMBL33 library represented with RDKit fingerprints (2048 bits) using the Russell-Rao (RR) index. Stratified methods use 10, 25, and 50 sampled molecules respectively. Three trials of a random selection of 50 molecules are shown.

**Figure 14:**
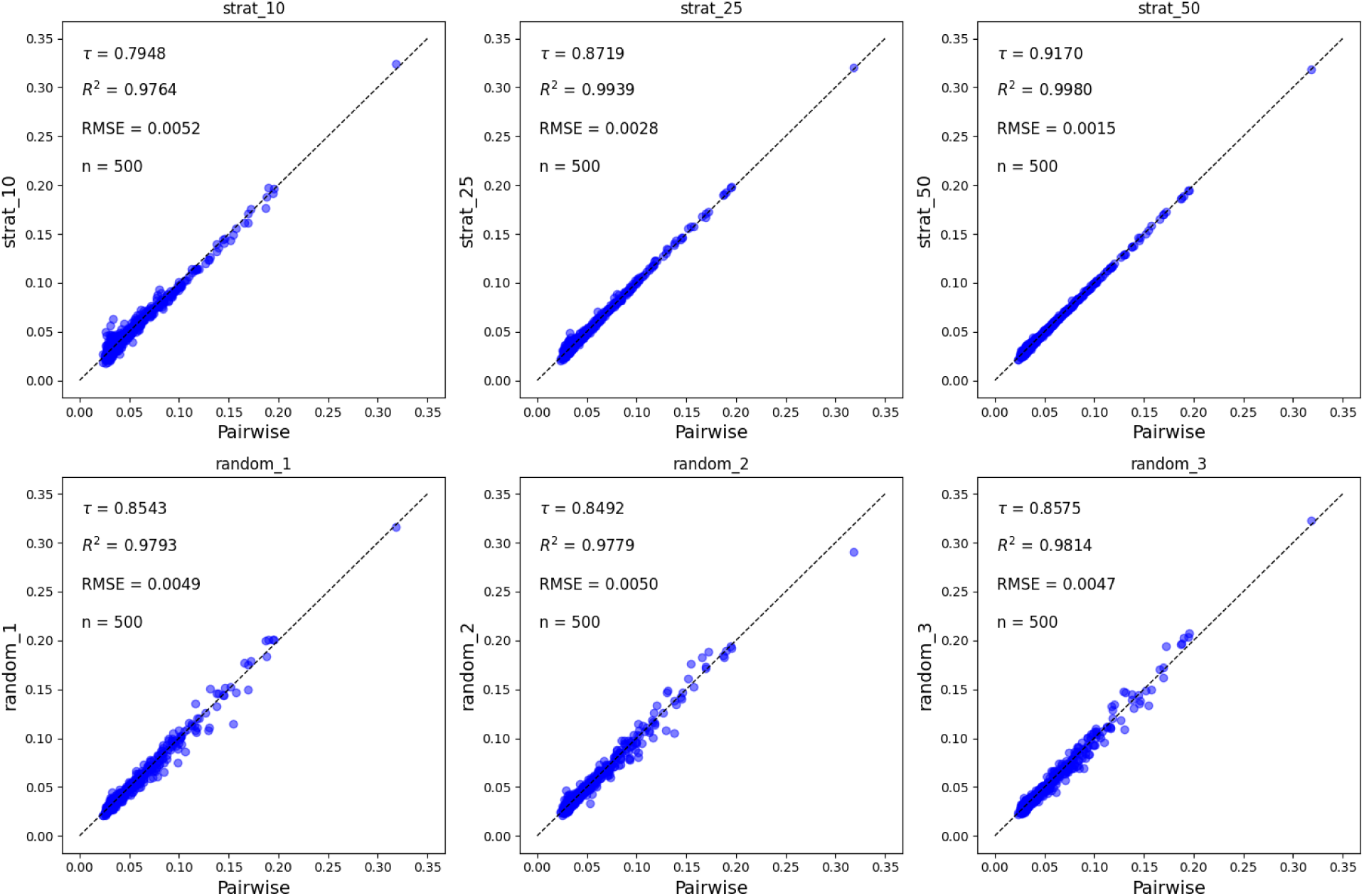
Comparison of the pairwise standard deviation of the similarity values and sampling approaches for 500 randomly selected subsets (sizes between 1000-5000) of the ChEMBL33 library represented with RDKit fingerprints (2048 bits) using the Sokal-Michener (SM) index. Stratified methods use 10, 25, and 50 sampled molecules respectively. Three trials of a random selection of 50 molecules are shown.

